# Thermal fluctuations assist high-speed rotation of the bacterial flagellar motor at low load

**DOI:** 10.64898/2026.09.09.750262

**Authors:** Shuichi Nakamura, Yusuke V. Morimoto, Nobunori Kami-ike, Tohru Minamino, Keiichi Namba

**Author notes:** Correspondence and requests for materials should be addressed to S.N., T.M. or K.N.

## Abstract

The bacterial flagellar motor is a proton-driven rotary nanomachine that converts ion flow into mechanical motion. Proton-coupled rotation of the MotA_5_-MotB_2_ stator complex generates torque, yet how this process drives high-speed rotation remains unclear. Here we use single-molecule nanophotometry to resolve stepwise rotation under low-load conditions. Lowering intracellular pH selectively prolongs dwell times without affecting step durations, indicating that proton dissociation triggers torque generation. Structural analysis further suggests that the rotor disengages from the stator before completing the elementary step angle (∼11°), implying that the power stroke alone is insufficient. Notably, the rotational diffusion coefficient of stator-less motors closely matches that inferred from wild-type motors at low load. These findings support a model in which intrinsic thermal fluctuations compensate for the limited reach of the power stroke, enabling rapid rotation. Our results reveal a hybrid mechanism in which ion-driven conformational changes bias stochastic motion to achieve efficient energy transduction.

## Introduction

Many motile bacteria, such as *Escherichia coli* and *Salmonella*, swim in liquid environments and move on solid surfaces by rotating long helical filaments with flagellar motors embedded in the cytoplasmic membrane. The flagellar motor of *E. coli* and *Salmonella* converts an inward-directed proton (H^+^) flow into the mechanical work required for flagellar motor rotation. The flagellar motor consists of a rotor and up to a dozen force-generating units called stators. The rotor is composed of the MS-ring made of the transmembrane protein, FliF, and the cytoplasmic C-ring, which consists of the proteins, FliG, FliM, and FliN. MotA and MotB form a transmembrane H^+^ channel that acts as a force-generating stator unit. Electrostatic interactions between MotA and FliG generate the rotational force. Two highly conserved residues, Asp of MotB (Asp-33 in *Salmonella* MotB) and Pro of MotA (Pro-173 in *Salmonella* MotA) are located in the H^+^ channel and are directly involved in H^+^ translocation across the cytoplasmic membrane^1–3^. The rate of H^+^ release from the MotAB channel into the cytoplasm limits the rate of flagellar motor rotation^4,5^, but it remains unknown how the MotAB complex couples H^+^ translocation with torque generation.

The number of active stator units around a rotor responds to changes in external load. The flagellar motor accommodates about ten stator units when the motor operates under high load. In contrast, the number of active stator units decreases from ten to as few as one with a decrease in external load^6–9^. Interactions between MotA and FliG affect the assembly and disassembly dynamics of the MotAB complex in a load-dependent manner^10^, suggesting that the MotA–FliG interaction not only generates torque for motor rotation but also controls the load-dependent dissociation of the MotAB complex from the rotor. The dissociation rate of the stator unit from the motor is much faster at extremely low load than at high load because the bound lifetime of each stator unit becomes shorter^11^. Consistently, the maximum speed of the flagellar motor at extremely low load depends on the number of active stator units in the motor^12,13^. A theoretical model has predicted that a duty ratio is small when the motor operates at extremely low load, whereas the duty ratio becomes high enough for the motor to generate much larger torque at high load^14^. However, it remains unknown how the motor operates over such a wide range of external loads.

High-resolution cryo-electron microscopy (cryo-EM) structural analyses have revealed that the MotAB stator complex consists of 5 MotA subunits and 2 MotB subunits^15,16^. The five MotA subunits form a pentameric ring (MotA_5_), while two transmembrane helices of the MotB dimer form a coiled-coil (MotB_TM-CC_) that penetrates the central pore of the MotA_5_ ring. These structural features support the hypothesis that the MotA_5_ ring rotates around the MotB_TM-CC_ in response to H^+^ transfer through the channel. In this model, the stator unit functions as a small gear turning the large gear of the rotor. The rotation of the stator complex labelled with an enhanced yellow fluorescent protein variant has been directly observed in living cells by polarized photobleaching macroscopy^17^. The stator rotation mechanism is also supported by molecular dynamics simulation^18^. This torque-generation mechanism, described as a power stroke, accounts for the high duty ratio under high load but does not explain why the duty ratio decreases when the motor operates at low load. To address this issue, we established a laser back-scattering nanophotometry system coupled to a high-speed camera^18^, operating at 390 kHz, to resolve the stepping motion of the H^+^-driven *Salmonella* flagellar motor at high speed. Our single-molecule experiments, together with theoretical analyses based on the recent cryoEM structure of the MotAB-FliG complex suggest that motor rotation at low load, driven by a single stator unit, involves Brownian motion following the power stroke caused by the MotA-FliG interaction.

## Results

### Spatial resolution of nanophotometry

Flagellar rotation consists of a torque generation step and a dwell time. However, it was unable to precisely measure the step times because each step was too fast to resolve by a nanophotometry system with temporal resolution at a sub-millisecond level^20,21^. Therefore, we developed a nanophotometry system to observe the rotation of a 100-nm gold nanoparticle (AuNP) (Extended Data Fig. 1a). The standard deviation in the position of an AuNP particle attached to the surface of a cover slip determined for every 2.56 μs image frame demonstrated that the accuracy of the determination of the AuNP particle position is better than 1 nm (Extended Data Fig. 1b). For a circular trajectory of rotating AuNP at a radius of 100-nm, for instance, our nanophotometry system is capable of determining its angular position to a 0.6-degree resolution or better.

### Direct observation of steps in high-speed rotation at low load

When the *Salmonella* flagellar motor operates at a load close to zero, large speed fluctuations and long pausing events are frequently observed. These fluctuations are suppressed by an increase in external load^11^. To observe fine steps generated by the *Salmonella* flagellar motor operating at low load, we attached a 100-nm gold particle onto a partially sheared sticky filament of the *Salmonella* Δ*cheA*-*cheZ* strain^5^, which rotates its flagella exclusively counterclockwise (CCW) due to deletion of chemotaxis genes. The probe movement was recorded at a frame rate of 390 kHz, and the change in the angular position of the probe was determined as it moved along its circular trajectory with a radius ranging from 100 to 150 nm (Fig. 1a, inset). Four angle-time traces of each revolution of a motor rotating CCW (CCW-locked wild-type motor) at speeds from 220 to 330 Hz are shown in Fig. 1a as typical examples (see also Movie 1). Steps and dwells were clearly observed (Fig. 1b and Extended Data Fig. 2). Steps with durations of several dozen microseconds were seen for many data points (Fig. 1b, inset). Backward steps were also observed occasionally (Extended Data Fig. 2).

**Fig. 1.**
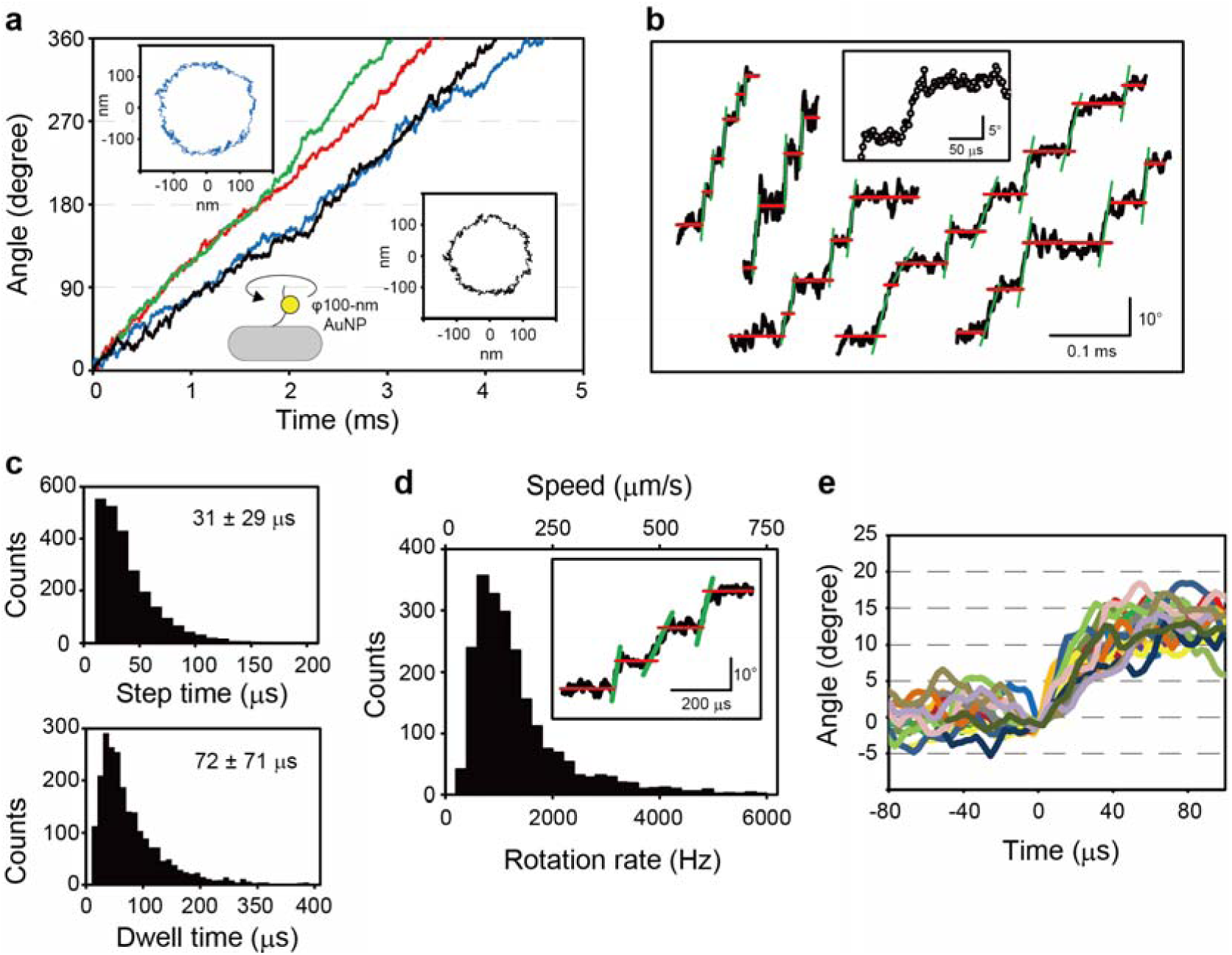
Stepping rotation. **(a)** A circular trajectory of a 100 nm AuNP attached to the sheared sticky flagellar filament of the CCW-loacked *Salmonella* strain MM3076iC rotating at around 300 Hz (inset) and its angular displacements plotted against time (black line). Data from four motors are shown in black, green, red, and blue. **(b)** Examples of steps and dwells detected in the angular displacements of CCW motor rotation. Black traces are raw data, and red and green lines are those automatically fitted to dwells and steps using the step-finding program (see *Methods*). A part of the traces expanded in the inset shows that demonstrates that the step period contains enough data points to measure its duration and slope. **(c)** Step time and dwell time distributions of the CCW-locked motor measured in motility medium, pH 7.0. **(d)** The distribution of step speeds calculated from slopes fitted to steps with the automated step finder (green lines shown in the inset). Step speeds are on two scales: The lower label is the rotational velocity (Hz), and the upper label is the linear velocity assuming the radius of the C ring to be 20 nm. **(e)** Selected steps for a single motor are superposed to show the fluctuation in step speeds.

The step time is defined as the duration during which the rotor is rotating as a result of the interaction between the stator and rotor, whereas the dwell time is the time period of the rotor bound to the stator. To understand the elementary process of the torque generation cycle of the flagellar motor, we used an automated step-finding program to determine both step and dwell times (see *Methods*). A histogram of the step times showed a single exponential decay, while that of the dwell time showed a double exponential distribution (Fig. 1c). The average step and dwell times were 30.6 ± 29.1 μs and 72.2 ± 71.0 μs, respectively. The double exponential distribution of the dwell times suggests that two reactions may occur during a dwell.

We next fitted a linear line to the step between two consecutive dwells (Fig. 1d, inset), determined the speed of each step from the slope, and made a histogram of speeds to determine the speed distribution (Fig. 1d). The average step speed was 1,626 ± 3,316 Hz (n = 2,290 CCW steps). Since 34 FliG subunits are located at the top of the C-ring and interact with the stator at a radius of about 20 nm, this frequency corresponds to approximately 200 μm/s as a linear speed along the circumference of the C-ring. The speed distribution was broad at high speed, extending beyond 4,000 Hz (∼ 500 μm/s) (Fig. 1d), with a few steps showing a speed as fast as 5,000 Hz. Thus, the rotation speed during the stepping events is more than 10 times higher than the average net speed. Superposition of many step traces shows a wide range of slopes (Fig. 1e). This contrasts with the relatively constant step rotation speed of about 850 Hz observed for the γ subunit of F_1_-ATPase^22,23^, which is driven by a conformational change of the α and β subunits upon binding and hydrolysis of ATP^24^. Therefore, we suggest that the step process of a flagellar motor operating at low load involves a stochastic event in addition to a predictable kinetic event.

### Effect of intracellular pH changes on step and dwell times

A decrease in intracellular pH by lowering external pH in the presence of a weak membrane-permeable acid markedly reduces the maximum rotation rate of the flagellar motor at low load. However, it does not affect the total proton motive force across the cytoplasmic membrane, suggesting that H^+^ dissociation from the H^+^ channel into the cytoplasm is the rate-limiting step of the torque-generation cycle at low load^4,5^. To determine which elementary process is affected by lowering cytoplasmic pH, we measured both step and dwell times at an external pH value of 7.0 or 6.5 in the presence of 5 mM potassium benzoate. In agreement with a previous report^5^, the average net speed of the CCW-locked wild-type flagellar motor was reduced with a decrease in intracellular pH from 7.5 to 6.9 (Fig. 2a and Extended Data Fig. 3a). However, the step times were unaffected, and their distribution profile was almost similar to that for the motor rotating at external pH of 7.0 in the absence of potassium benzoate (Fig. 2b, left panels and Fig. 2c, left panel). In contrast, the distribution of dwell times broadened, indicating that the reduction in the average net speed at lower intracellular pH is a consequence of a longer dwell time (Fig. 2b, right panels and Extended Data Figs. 2 right panels and 3b). The average dwell time increased from 70 to 120 μs by lowering internal pH from 7.5 to 6.9, and the longest dwell time increased from around 300 to 600 μs. The distribution profiles of the dwell times were fitted by a double exponential with reaction rates of 30 ms^-1^ and 31 ms^-1^ at an external pH 7.0 in the absence of potassium benzoate. Both rate constants were reduced with a decrease in intracellular pH (Fig. 2c, right panel). Because the rotor is strongly bound to the stator during the dwell time, we suggest that the dissociation of H^+^ from the stator into the cytoplasm induces the dissociation of the rotor from the stator for step rotation.

**Fig. 2.**
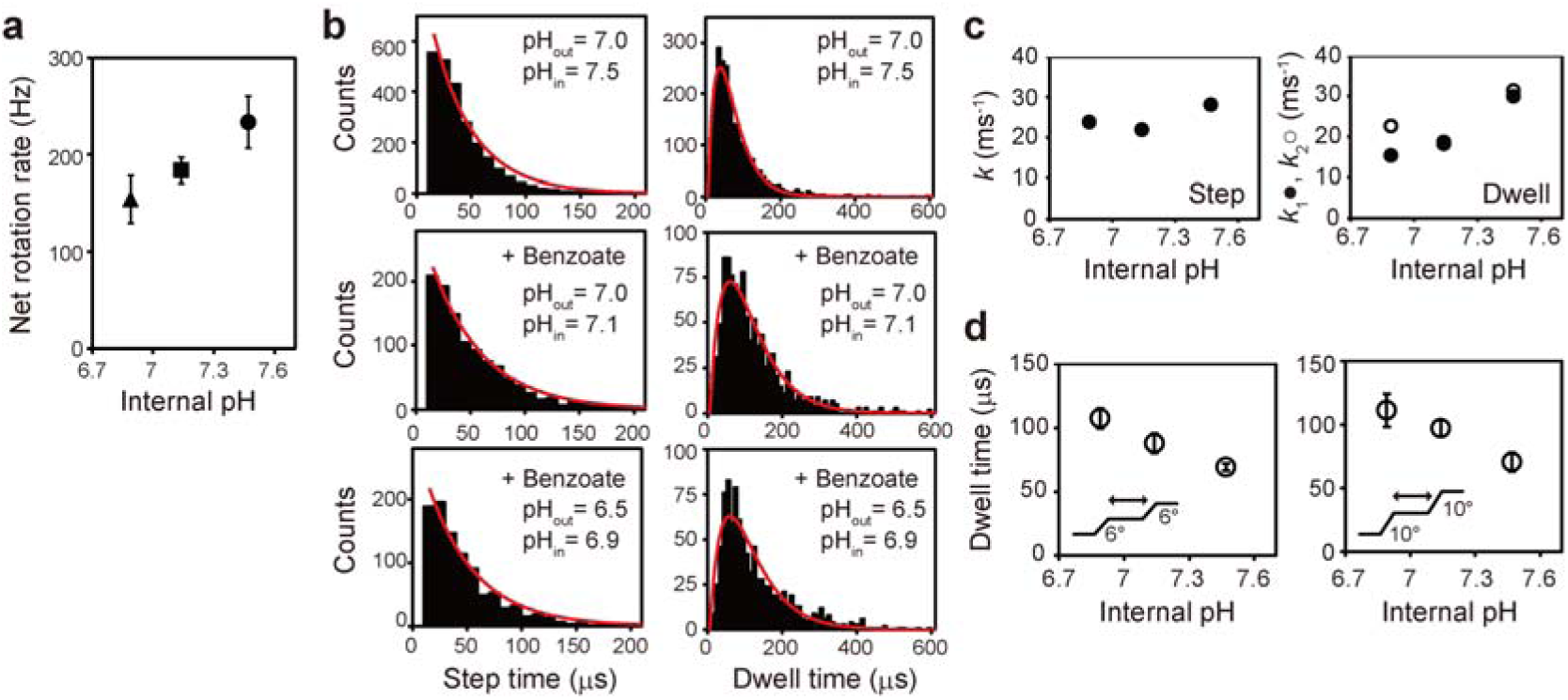
Effect of low intracellular pH on the step and dwell times. **(a)** Effect of internal pH changes on the net speed of the flagellar motor. Net rotation rates decreased when the external pH was lowered in the presence of 5 mM potassium benzoate. Average values of data measured in motility medium (pH 7.0) without benzoate (circle), in motility medium (pH 7.0) containing benzoate (square), and in motility medium (pH 6.5) containing benzoate (triangle). Error bars show the standard error. The internal pH was measured using pHluorin(M153R), which is a pH indicator protein. **(b)** Histograms of step and dwell times. The histograms obtained in motility buffer without benzoate (upper panels) were the same as in Fig. 1c. Distributions of step and dwell times were fitted by a single exponential, *A*×exp(−*kt*), and double exponential, *A*×[exp(−*k*_1_*t*)−exp(−*k*_2_*t*)], respectively (red lines). **(c)** Effect of intracellular pH on rate constants of step (left) and dwell (right). These two rate constants were obtained as fitting parameters in b. **(d)** Effect of intracellular pH on dwell times. Data of dwell times were grouped according to step angles determined by multi-Gaussian fitting to distributions of step angles (Extended Data Fig. 4). The step angles and the position of the dwell period in each group are depicted in insets. The average values and standard errors are shown.

The histogram of step angles showed a wide distribution (Extended Data Fig. 4). When it was fitted by multi-Gaussian functions, three distinct peaks appeared at around 6°, 10°, and 13° (Extended Data Table 1). The number of functional stator units ranges from one to five when the flagellar motor operates at low load^25^. Because the FliG-ring has a 34-fold rotational symmetry in the *Salmonella* flagellar motor^26–28^,10° steps should reflect an actual interaction between MotA and FliG (360°/34 FliG subunits = 10.6°) when the motor is rotated by a single stator unit.

The number of functional stator units driving the rotor measured in the present study was estimated to be one or two from the net speeds determined for each revolution (Extended Data Fig. 5). Because the number of active stator units fluctuates at extremely low load^11,13^, frequent changes in the step angle may be a consequence of changes in the number of active stator units at low load. Assuming that the number of functional stator units is variable, smaller steps might be observed upon incorporation of additional stator units. Therefore, in addition to the 10° steps, the 6° steps may reflect an elementary torque generation process. The 13° steps have been observed when the motor rotates at a few Hz^20,21^. Because the LP-ring that acts as a molecular bushing for high-speed rotation of the rod that acts as a drive shaft of the flagellar motor has a 26-fold rotational symmetry^27^, we assume that the 13° step angle may reflect an interaction between the rod and the LP-ring.

We classified dwell times according to the step angle (Fig. 2d, insets) and found that lowering the internal pH prolonged dwell times at all step angles (Fig. 2d). This suggests that dissociation of H^+^ from the MotAB channel into the cytoplasm is a critical step for high-speed motor rotation under low load. Although it was difficult to quantify the kinetics of the dwell times, the observed double exponential distribution (Fig. 1c and Fig. 2b right panels) suggests that each dwell comprises at least two distinct reaction processes, involving the protonation-deprotonation cycle of Asp-33 in MotB coupled with conformational changes within the H^+^ channel.

### Estimation of rotation angle by direct interaction between MotA and FliG

MotB_TM-CC_ penetrates the central pore of the MotA_5_ ring. As a result, two distinct H^+^ translocation pathways involving the Asp-33 residues of MotB are created. The MotB_TM-CC_ also acts as an axle for the MotA_5_ ring that rotates around it like a wheel. The movement of H^+^ through the channel induces unidirectional rotation of the MotA_5_ ring relative to this axle, allowing interactions between MotA and FliG to spin the rotor. Thus, all five MotA subunits are involved in torque generation, and the power stroke is generated by the rotation of the MotA_5_ ring relative to MotB_TM-CC_. The flow of H^+^ through the channels presumably causes the MotA_5_ ring to rotate only in the CW direction^15,16^. The flagellar motor can rotate both CCW and CW without changing the direction of the H^+^ flow through the channel. Thus, directional switching must depend upon conformational changes in the C-ring that affect the way the rotor interacts with the MotA_5_ ring rotating in the CW direction. Consequently, when the MotA_5_ ring undergoes a 36° rotation, the rotor rotates forward by about 11° according to its conformational state.

We examined whether the rotor ring can rotate up to 11° while remaining bound to the rotating MotA_5_ ring using the atomic models of the *Clostridium sporogenes* MotAB complex with the C-terminal domain of FliG (FliG_C_) (PDB ID: 8UCS) and the CCW-state C-ring of *Salmonella enterica* (PDB ID: 8UOX)^29^. FliG_C_ from the 8UCS structure was superimposed onto the equivalent coordinate of the 8UOX structure (Fig. 3a), revealing tight interactions between MotA and FliG_C_ (Fig. 3b). However, critical binding pairs (MotA Arg89 and Arg93 with FliG Glu292, and MotA Glu100 with FliG Arg285) are disrupted by a 2° CCW rotation of the rotor ring, accompanied by a 6° CW rotation of the stator, thereby releasing FliGc from MotA (Fig. 3c,d). These structural analyses suggest that the rotor rotates at most ∼2° through the MotA-FliG_C_ interactions as the MotA_5_ ring rotates CW. Further rotation of the MotA_5_ ring, in conjugation with the rotor, results in the disengagement of FliG_C_ from MotA (Extended Data Fig. 6, Movie 2, and Movie 3). Thus, the power stroke generated by the MotA-FliG_C_ interaction alone is insufficient to rotate the rotor by ∼ 11°. Under conditions where the motor operates with a single stator unit at low load, the remaining angle of rotation must therefore be completed in a stator-independent manner.

**Fig. 3.**
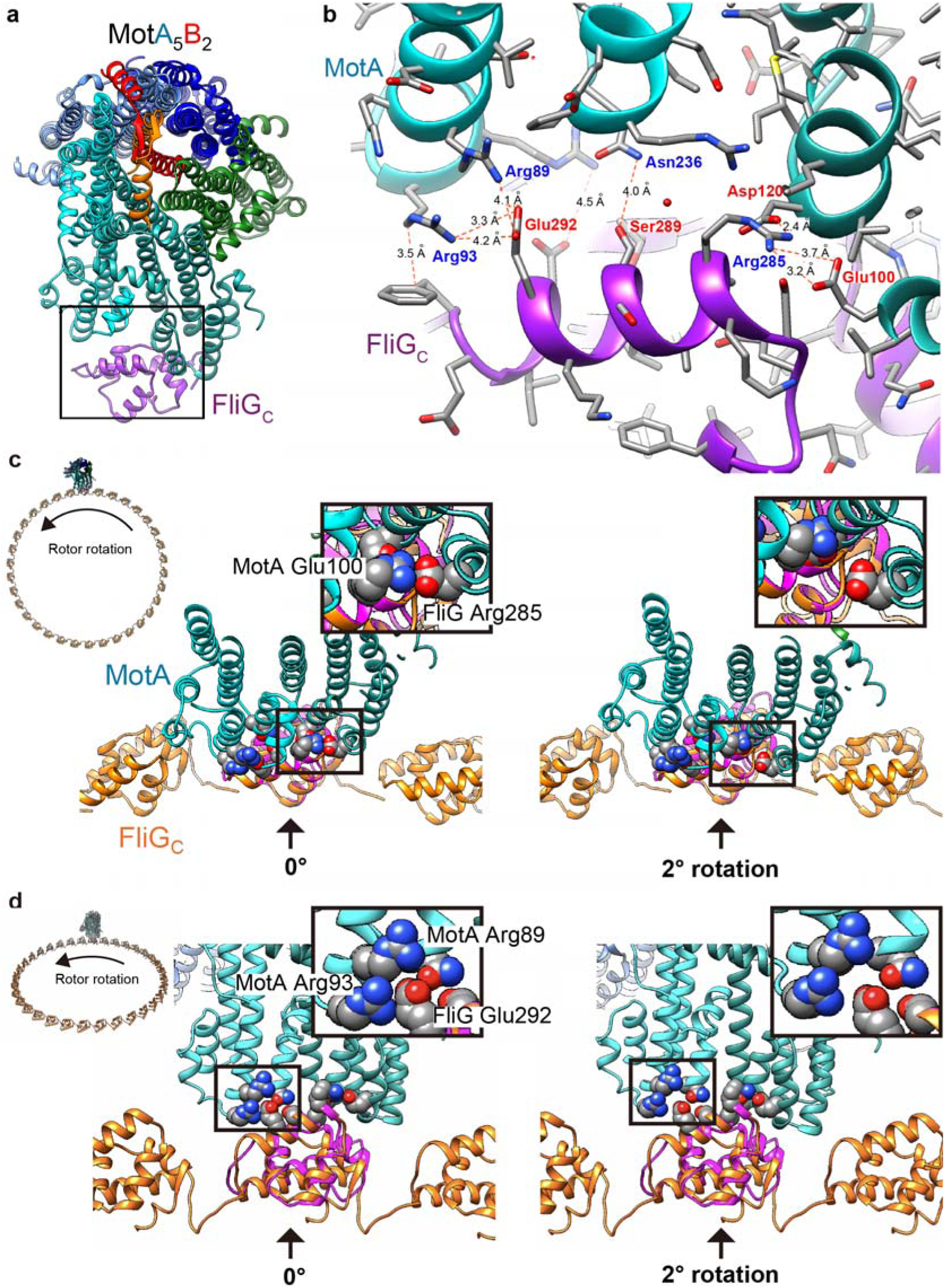
Structural limits of the MotA-FliG interaction. **(a)** The relatively tight bonding interactions of MotA and FliG_C_ (*C. sporogenes*, 8UCS) in the cryoEM structure of the complex of MotA_5_B_2_ and FliG_C_ by Johnson *et al*.^29^. **(b)** Enlarged view of the area marked with a black rectangle in ***a***. **(c)** The FliG_C_ model in the MotAB-FliG_C_ complex structure is superimposed on one of 34 FliG_C_ subunits in the C ring of the flagellar basal body structure (*Salmonella enterica*, 8UOX)^29^ and viewed along the rotation axis of the basal body (viewed angle is shown in the inset, top left). Arrows indicate 2° rotation of a FliG subunit. The areas enclosed with black rectangles are enlarged in the top right, showing the binding and dissociation of MotA-Glu100 and FliG-Arg285 at 0° and 2°, respectively. See Supplementary Movies 2 (entire ring) and 3 (enlarged view as shown in ***c***) for the time-series changes in FliG-MotA interactions accompanying the mutually counter-rotating rotor and stator. **(d)** FliG_C_-MotA interaction viewed from different angles (shown in the inset, top left). The black-boxed regions indicate the dissociation of FliG-Glu292 from MotA- Arg89 and MotA-Arg93 upon 2° rotation of the FliG ring.

### Rotational diffusion coefficient of the flagellar motor with or without the stator units

Based on the structural analyses of the MotA-FliGc interaction, we assumed the contribution of Brownian motion to the stepping movement of the flagellar motor. To verify this hypothesis, we analyzed the diffusivity of stepping movements detected in the CCW-locked wild-type motor and compared it to that of the stator-deficient (ΔMotAB) motor, where free rotational diffusion has been observed^11,30^.

The radius of rotation of the gold probe attached to the ΔMotAB motor was ∼100 nm, similar to that observed for the wild-type motor (Fig. 4a). We analyzed the mean square displacement (MSD) of the ΔMotAB motor and found that the rotational diffusion coefficient (*D*_rot_) was ∼270 rad^2^/s (Figs. 4b and 4c). Diffusion analysis of the CCW-locked wild-type motor was performed using the first-passage time distribution (FPTD), which is a distribution of times required for the diffusion-controlled displacement of an arbitrary distance (*ϕ*_step_)^31^. Assuming a reflection boundary at *ϕ* = 0 and an absorption at *ϕ* = *ϕ*_step_ (= 6° or 10°), the FPTD fitted well to the measured step-time distributions regardless of pH and step angle (Fig. 4d). The values of *D*_rot_ obtained as the best-fit for the wild-type distributions were 200-500 rad^2^/s, a range that is comparable to the 270 rad^2^/s of the ΔMotAB motor (Fig. 4c). This equivalence suggests that rotational Brownian motion is a major contributor to the step rotation of the flagellar motor operating at low load.

**Fig. 4.**
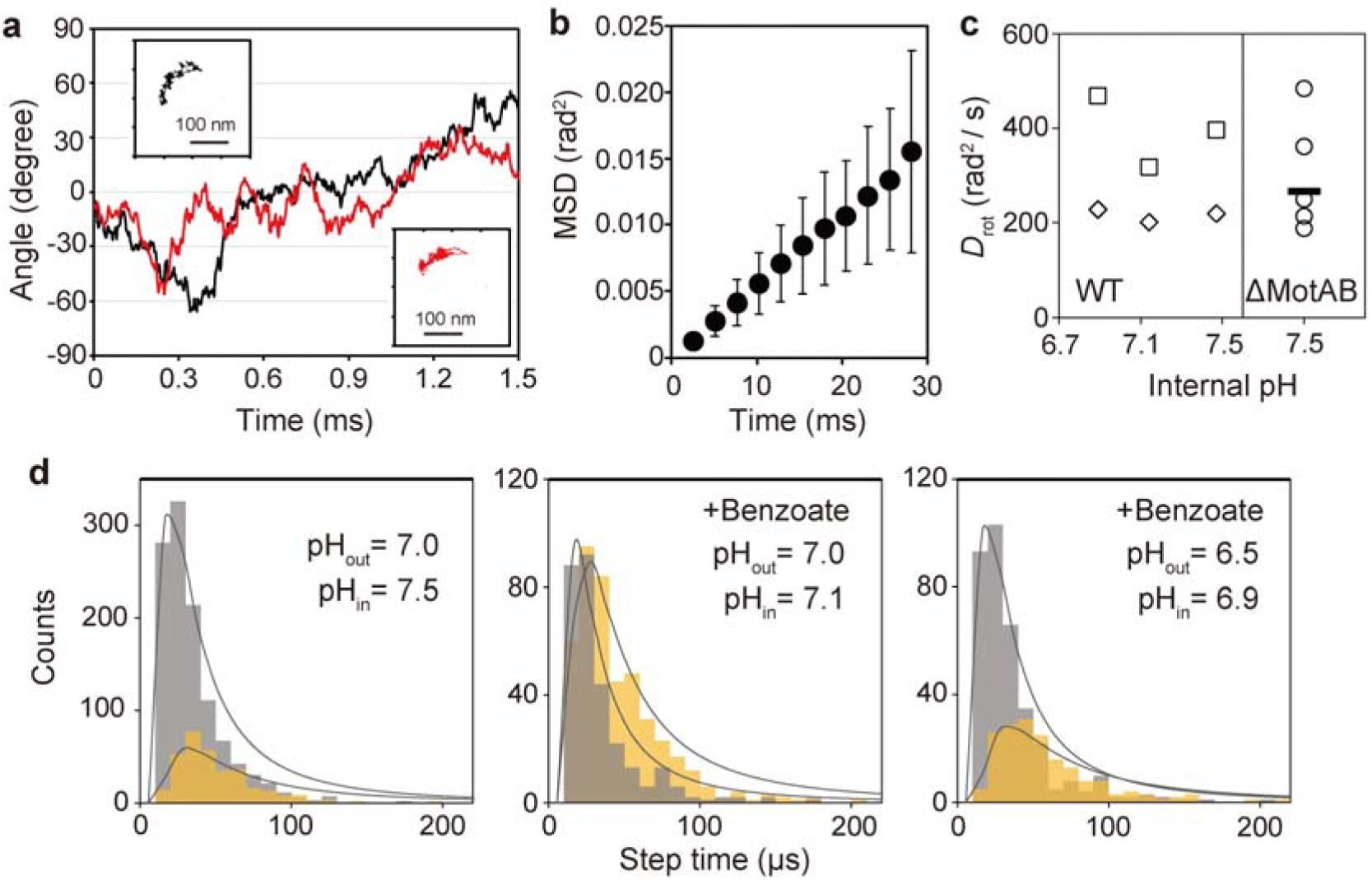
Diffusion analysis. **(a)** Angular displacement data for the flagellar motor without stator units. Two data sets are shown in black and red as examples. The trajectories of AuNPs used for the angular displacement data are shown in the insets. **(b)** MSD analysis for the stator-less mutant motor. Average values and standard deviations from 5 motors are shown. **(c)** Values of the rotational diffusion coefficient *D*_rot_ obtained by MSD and FPTD analyses (shown in ***d***) of the CCW-locked wild-type (WT) and stator-less mutant (ΔMotAB) motors. For the ΔMotAB motor, data for 5 different motors and the average value are shown as circles and a horizontal bar, respectively. Squares and diamonds represent the data for 6° and 10°steps seen in the WT motor, respectively. **(d)** FPTD of steps found in CCW-locked wild-type motors measured in motility medium (pH 7.0) without benzoate, in motility medium (pH 7.0) with 5 mM potassium benzoate, and in motility medium (pH 6.5) with 5 mM potassium benzoate. Data were classified into three step angle groups as shown in Fig. 2d. Grey and orange histograms indicate the time distribution for 6° and 10° steps, respectively. Gray lines show the results of fitting by a theoretical FPTD^31^: *A* [*L*_step_/(2(π*D*_rot_*t*^3^)^0.5^)] [exp(−*L*_step_/4*D*_rot_*t*)−exp(−*L*_step_^2^/*D*_rot_*t*)], where *A* is a constant, *L*_step_ is the step angle, and *D*_rot_ is shown in ***c***.

The automated step finding program detected “dwells” even in the ΔMotAB motor (Extended Data Fig. 7a). However, the dwells in the wild-type motor were clearly different from those in the ΔMotAB motor (Extended Data Fig. 7b): the variance in the probe position in the dwell time of the wild-type motor was substantially less than that in the ΔMotAB motor (Extended Data Fig. 7c). Therefore, we suggest that the dwell time observed in the CCW-locked wild-type motor reflects the time when the stator unit remains firmly attached to the rotor, whereas the dwell times seen in the ΔMotAB motor might be due to transient pauses caused by friction between the motor and the cytoplasmic membrane and/or between the rod and the LP-ring complex.

### Thermal fluctuations overcome the structural limits of the MotA-FliG interaction

The torque versus speed relationship of the *Salmonella* flagellar motor has been extensively investigated^5,32–34^. The maximum torque produced by the *Salmonella* flagellar motor is estimated to be about 2,000 pN nm^5^. The flagellar motor can accommodate ten stator units around a rotor when the motor operates at high load^10^. The average rotation rate of the *Salmonella* flagellar motor was about 250 Hz when the partially sheared filament was labeled with a 100-nm nanoparticle^5^. The flagellar bundle of a *Salmonella* cell swimming in a low-viscosity liquid medium rotates at 100-170 Hz^35^, which must be the rotation rate of each single flagellum in the bundle. Based on that the combination of a power stroke by the MotA-FliG interaction with Brownian motion of a rotor free from the stator likely drives motor rotation at low load, we attempted to explain the known motor characteristics theoretically as follows (a hypothetical reaction scheme is shown in Fig. 5a; values of parameters are listed in Extended Data Table 2):

(1) The dwell time, *t*_dwell_, is determined by the rate constant of H^+^ dissociation from the stator unit into the cytoplasm, *k*_H_^Cyto^ and the detach rate of the rotor from the stator, *k*_off_^R^: *t*_dwell_ = 1/(*n*_s_ × *k*_H_^Cyto^) + 1/(*k*_off_^R^ / *n*_s_), where *n*_s_ is the number of active stator units around a rotor. The dissociation rate of H^+^ apparently increases with increasing *n*_s_ whereas the probability of rotor detachment decreases with increasing *n*_s_, because all stator units must detach from the rotor at the same time for the rotor to undergo rotational diffusion in the subsequent step time.
(2) The step size *ϕ* depends on *n*_s_ by *ϕ* = *ϕ*_0_/*n*_s_, where *ϕ*_0_ is the step size when the motor is operated by a single stator unit; we here assume *ϕ*_0_ ∼11° (360°/ 34 FliG subunits^28^).
(3) The potential for power stroke *U*(*θ*) = −*N*_ps_*θ*_ps_, where *N*_ps_ is a constant torque, and *θ*_ps_ is the angle of the rotation driven by the power stroke. *θ*_ps_ = 2° was assumed by atomic model-based verification of the geometrical limit of MotA-FliG interaction (Fig. 3). The free energy of H^+^ (∼7*kT* at external pH = 7.0, internal pH = 7.4, Δ*φ* = 150 mV, and 23°C)^5^ is utilized to generate the structural change changes in the stator unit that drive the power stroke (Fig. 5b). The rotation of the rotor driven by the power stroke of a single stator unit occurs in *t*_ps_ ∼ *θ*_ps_/*ω*_ps_ = *θ*_ps_/(*N*_ps_/*γ*), where *ω*_ps_ is the drift angular velocity and *γ* is the drag coefficient.
(4) The rotor moves through the remaining angle *θ*_diff_ (= *ϕ* − *θ*_ps_) by free diffusion of the rotor with a duration of *t*_diff_ ≈ *θ*_diff_^2^/2*D*_rot_ = *θ*_diff_^2^/(2*k*_B_*T*/*γ*). Therefore, *t*_rot_ = *t*_ps_ + *t*_diff_.
(5) The step time, *t*_step_, is determined by either a rotation time, *t*_rot_, or a waiting time for the binding of H^+^ to the Asp-33 residue in the H^+^ half-channel open to the periplasm, *t*_H_^Peri^. If *t*_rot_ < *t*_H_^Peri^, the stator unit is not able to bind to the rotor until it is protonated, and therefore *t*_step_ = *t*_H_^Peri^. If *t*_rot_ > *t*_H_^Peri^, the stator unit waits until the rotor reaches a catchable position, and therefore *t*_step_ = *t*_rot_. The time required for H^+^-binding to a Asp33 residue exposed to the periplasm is determined by *t*_H_^Peri^ = 1 / (*n*_s_ × *k*_H_^Peri^), where *k*_H_^Peri^ is the H^+^-association rate in the periplasm.
(6) The net rotation rate *ω*/2π is given by 1/(*t*_dwell_ + *t*_step_), and the motor torque is equal to *γ* × *ω*.

**Fig. 5.**
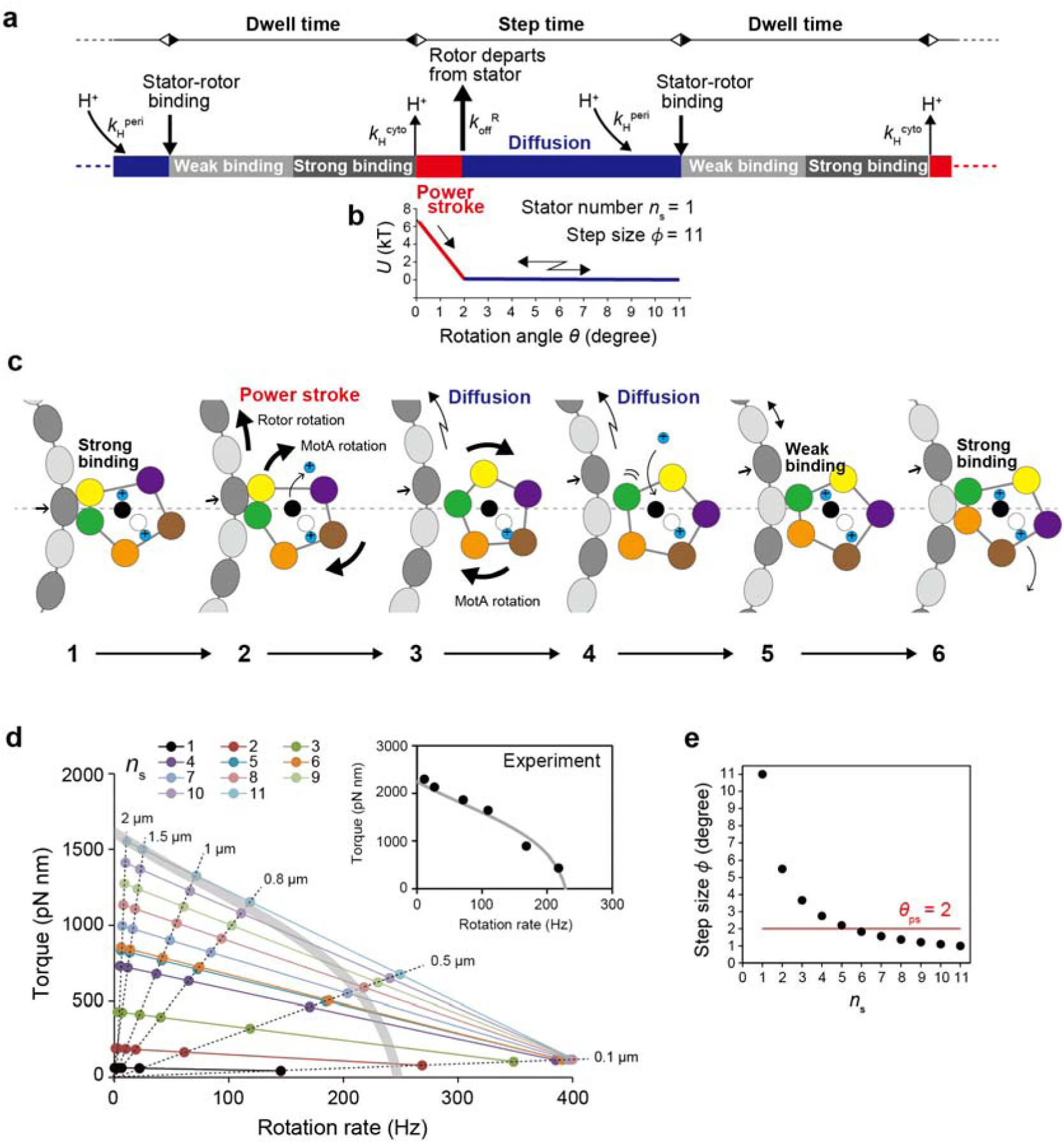
Rotation model of the flagellar motor. (a) Hypothesized scheme of the flagellar motor rotation. (b) The potential landscape in the stepping period when the rotor is rotated by a single stator unit (*n*_s_ = 1). The rotor is rotated 2° (= θ_ps_, see Fig.3 and the main text) by power stroke immediately after detachment from the stator. (c) Modelling the coupling of protonation/deprotonation with the rotation of the stator and rotor. (d) The torque-speed relationship predicted by the model. 2.0 μm, 1.0 μm, 0.8 μm, 0.5 μm, and 0.1 μm beads were assumed for the calculation. The number of active stator units are indicated by dots with different colours. The thick grey line is an approximate torque-speed relationship, arbitrarily written to be consistent with the experimental data (inset, drawn referring to Nakamura *et al*.^5^). (e) Dependence of the step size on the number of active stator units. The red line indicate the threshold angle for power stroke, estimated from the structural limits of the MotA-FliG engagement.

Fig. 5c depicts structurally compatible models for the coupling of the H^+^ flux through the H^+^ channel with rotations of the MotA_5_ ring and a rotor. Based on the stator-unit rotation model proposed by Santiveri *et al*.^15^, we assumed that at least one of the two MotB subunits is always protonated. When both H^+^ channels are occupied by H^+^, two MotA subunits in the ring (yellow and green) strongly bind to a single FliG subunit (1), rendering the MotA_5_ ring asymmetric. The dissociation of one H^+^ into the cytoplasm induces rotation of the MotA_5_ ring and drives FliG by a power stroke (2). The MotA-FliG interaction disengages after ∼2° rotation (3), allowing the FliG ring to diffuse. Given that the conformational symmetry of the MotA_5_ ring without the interaction with FliG is barely distorted by the protonation of the critical proton binding site Asp-32 in MotB^15^, the MotA_5_ ring restores to a symmetric conformation (4), and one MotA subunit (green) weakly binds to the nearest FliG subunit (5). The binding of another MotA subunit (orange) to the FliG subunit distorts the MotA_5_ ring into an asymmetric conformation and provides the strong-binding state (6). Comparing the state 1 and the state 6, the MotA_5_ ring appears to rotate ∼72°, but it is mostly attributed to the distortion and restoration of the MotA_5_-ring conformation, and the movement of one MotA subunit directly involved in catching FliG (green) is approximately half of the rotation of the other MotA subunits. This model also explains that lowering the intracellular pH slows down the H^+^ dissociation from the H^+^ channel into the cytoplasm, thereby prolonging the dwell time (Fig. 2d).

This kinetic model reproduced the previous experimental data of torque-speed curve (Fig. 5d)^5,10,32–34,36^. The model suggests that more than 50% of the step angle resulted from free diffusion without a power stroke in the motor driven by one or two stator units at low load. Because diffusion is inversely proportional to external load, the diffusion-dependent movement will be effective at low load but not at high load. On the other hand, the contribution of the power stroke is increased with *n*_s_ obeying the relation *ϕ* = *ϕ*_0_/*n*_s_ (Fig. 5e). Since *n*_s_ increases with *γ* ^6,7^, the flagellar motor is predominantly driven by the power stroke at high load. Because the bound lifetime of each active stator unit is much shorter at low load than at high load^11^, such changes in the strength of the MotA-FliG interaction may be responsible for autonomous control of the number of active stator units in the motor in response to changes in external load.

## Discussion

The FliF–FliG deletion fusion mutation in *Salmonella*, which deletes 56 residues from the C-terminus of FliF and 94 residues from the N-terminus of FliG and fuses them together, not only causes a strong CW switch bias but also inhibits high-speed rotation of the flagellar motor at low load. A single stator unit can rotate the FliF–FliG deletion fusion motor with a 1.0-μm bead attached at the same level as the wild-type motor, indicating that the FliF-FliG deletion fusion mutation does not affect the energy coupling efficiency of the flagellar motor. Because the rotational speed of the flagellar motor operating in the low-torque, high-speed regime depends on the mechanochemical reaction cycle of the motor, the FliF-FliG deletion fusion restricts conformational dynamics of the MotA_5_ ring coupled with H^+^ flow through the transmembrane H^+^ channel of the stator unit at low load. Extragenic suppressor mutations in FliG, FliM, or FliN in the C-ring of the FliF-FliG deletion fusion motor not only relieve the strong CW switch bias but also increase the maximum rotational speed of the deletion fusion motor significantly. These suppressor mutations are located at an interface between the C-ring proteins, suggesting that a change of intersubunit interactions between the C-ring proteins may be required for high-speed motor rotation as well as directional switching^36^. Because the duty ratio of the motor decreases as the external load decreases^11–13^, the operation mechanism of the flagellar motor may be different at low and high loads.

We showed stepping rotation with ∼11° intervals in the *Salmonella* flagellar motor driven by a single stator unit at low load, which is consistent with the rotational symmetry of FliG ring (∼360°/34)^28^. We identified the kinetic properties of each step, which are attributable to a power stroke mediated by direct MotA-FliG_C_ interactions coupled with proton translocation. A superposition of the MotAB complex with the FliG_C_ domain on the equivalent coordinate of the C-ring revealed that the maximum rotation angle of the rotor achieved by the binding of the rotating MotA_5_ ring with FliG_C_ is approximately 2°. Based on the structural limitation that power stroke can contribute to rotation, we found that thermal fluctuations allows the rotor to rotate the remaining angle.

In order to allocate minimum functionality as a motion mechanism to a microscopic system, it is necessary to incorporate some factors that bias Brownian motion in some form. Biased Brownian motion has been adopted to explain stochastic but processive movement of a single-headed kinesin, KIF1A, toward the plus-end of microtubule (MT)^37^. KIF1A associated with ADP weakly binds to the MT and shows bidirectional one-dimensional Brownian diffusion along the MT. However, the movement becomes biased in the presence of ATP^38^, suggesting that going through the strong binding state by ATP hydrolysis is the key process for the biased directional movement. Dynein is a motor protein toward the minus-end of the MT in response to ATP hydrolysis. Mutations of two α-tubulin residues that form salt bridges with the MT-binding domain (MTBD) of dynein result in the loss of strong MTBD-MT binding, loss of activation of the dynein ATPase activity, and loss of directional movement^39^. MTs with mutations in either of these two α-tubulin residues show one-dimensional Brownian diffusion in the gliding assay on dynein-coated glass surface^39^. A cryo-EM structure of F-actin fully decorated by myosin heads of skeletal muscle provides a structural clue as to how directionally asymmetric dissociation of F-actin and myosin can be achieved by their asymmetric interaction in the weak-binding state and suggests a Brownian ratchet mechanism for the actomyosin motor during muscle contraction^40^. These results indicate that switching between the strong and weak binding states in response to ATP hydrolysis is key to the biased Brownian movement. In addition to biomolecular motors, grain boundary asymmetry in inorganic polycrystals has been shown to drive Brownian ratchet-like crystal growth^41^. In the flagellar motor, the flow of H^+^ through the two H^+^ channels in the MotAB complex rotate the MotA_5_ ring only in the CW direction due to steric hindrance between MotA and MotB. The biased rotation of the MotA_5_ ring generates a power stroke to push the rotor CCW (Fig. 5c, 2) and induces directionally asymmetric dissociation of the rotor from the stator, followed by the rotational diffusion biased towards CCW. Thus, directionally asymmetric dissociation of two-component proteins from their strong binding state appears to be a common mechanism to bias Brownian motion to produce unidirectional movement of motor proteins.

The number of stator units incorporated into motors of bundled flagella in swimming bacteria is likely enough to be driven only by the power stroke (*n*_s_ ∼5 in *E. coli*^42^), precluding the expectation of a significant contribution from Brownian motion. Why has the rotational mechanism of the flagellar motor by Brownian motion been retained in the evolutionary process? The flagellar motor can rotate even at the PMF as low as-30 mV, at which the free energy associated with proton translocation is close to that of thermal fluctuation (∼26 meV at 23°C)^43^. Since the dissociation of the stator units is facilitated by the PMF disruption^44^, the number of docked stator units would be decreased drastically in such low PMF. The Brownian motion-associated mechanism may allow bacteria to maintain locomotion under very low energy conditions, where the number of stators to be incorporated into the motor is limited. This intrinsic mechanism, although inefficient, but operates in a stochastic manner at low energy levels, thereby enhancing the robustness for bacterial survival and aiding in the acquisition of their ecological niche.

## Methods

### Bacterial strains and media

*Salmonella enterica* serovar Typhimurium strains MM3076iC [Δ(*cheA*-*cheZ*), *fliC*(Δ204–292)]^5^ and YSC2123 [Δ(*cheA*-*cheZ*), Δ*motA*-*motB*, *fliC*(Δ204–292)]^45^ were used in this study. The flagellar motor of the MM3076iC strain is intact but rotates exclusively CCW because the chemotaxis genes of the strain are deleted. The flagellar motor of the YSC2123 is deleted for the *motA* and *motB* genes and thus has a Mot^-^ phenotype. Both strains produce sticky flagellar filaments because of the *fliC*(Δ204–292) mutation, which allows gold nanoparticle (AuNP) to attach easily to the sheared flagellar filaments^8^. L-broth contains 1.0% (w/v) tryptone, 0.5 % (w/v) yeast extract, and 0.5% (w/v) NaCl. Motility medium contains 10 mM potassium phosphate, 0.1 mM EDTA, and 10 mM sodium lactate. The pH of the motility buffer was adjusted to the desired values by the addition of HCl or KOH in the presence or absence of 5 mM potassium benzoate.

### Sample preparation for bead assays

*Salmonella* cells were grown overnight in L-broth at 37`°`C with shaking. Overnight cultures were diluted 1:100 into fresh L-broth and incubated at 37`°`C for 3 h. The cells were collected and suspended in motility medium, and their filaments were sheared by passing cells through a 25G needle. The cell suspension was infused into a flow chamber and incubated for 20 min at 23`°`C. Cells floating in the chamber without attachment to the glass surface were washed away with motility buffer, and 100-nm AuNPs (BBI solutions) were added, followed by incubation for 5 min at 23`°`C. Unbound AuNPs were removed by washing with motility medium. To reduce the intracellular pH, cells were exposed to motility medium with or without 5 mM potassium benzoate at an external pH value of 6.5, 7.0, or 7.5. Intracellular pH of *Salmonella* cells was measured using a pH indicator protein, pHluorin(M153R)^46^ as described previously^5^. The diameter of the 100-nm AuNP was measured to be 117 ± 17 nm by electron microscopy.

### Optical system and observation of motor rotation

The back-scattering nanophotometry system was built on an inverted microscope (IX70, Olympus) as described by Sowa *et al.* 2010^19^, with some modifications. The specimen was placed on an ultra-stable stage with a two-dimensional piezo-micromanipulator (P-734, PI) that was used for adjusting the position of the AuNP to record its motion with an ultra-high-speed camera (Phantom V711, Vision Research Inc.). The 100 nm Au-NP attached to a sticky flagellar filament stub was visualized by illuminating with a He-Ne laser (30 mW, 05-KHP-991, Melles Griot), and the image of the AuNP produced by back-scattered light was observed through an oil immersion objective lens (100×, PlanApo, NA 1.4, Olympus). The camera can be operated at a maximum rate of 1.4 million frames per second for an imaging area of 128×16 pixels, but we operated it at 390 kHz (390,810 frames/s) to utilize a larger imaging area (128×64 pixels). The images of AuNP motion were thus recorded at intervals of 2.56 μs. The position of the AuNP was determined by fitting a two-dimensional (2D) Gaussian function to the peak intensity profile of the AuNP. The σ values of the Gaussian functions were around 100 nm. A wide dynamic range of the camera with 12 bits digital data recording (4096 grey scales) was important to determine the position of the AuNP with high precision. All measurements were carried out at 23`°`C.

The precision of the system in determining the position was estimated by imaging a 100-nm AuNP firmly attached to a cover slip. The X and Y positions of the AuNP were determined by 2D Gaussian fitting for each image every 2.56 μs, and the variations in their positions were analysed by fitting 1D Gaussian functions to the histograms (0.1 nm per bin) for the 392 X and Y positions recorded over 1.0 ms. The smallest 2σ of the Gaussian functions obtained at the highest intensity without sensor saturation was 0.82 nm in X and 0.6 nm in Y, indicating that the precision of this system is better than 1 nm.

### Rotation analysis

Elliptic trajectories of rotating AuNPs were corrected for the tilt of 3D trajectory plane to determine precisely the angular position on the circular trajectory in each image frame, as described previously^21^. The radius of rotation was determined from the elliptic parameters determined by fitting an ellipsoid to the trajectory. For a circular trajectory with a radius of 100-nm, our nanophotometry system is capable of determining the angular position at 0.6-degree resolution or better for each 2.56 μs image frame.

Steps and dwells were automatically identified by a step-finding algorism developed in this study. The angle positions plotted against time were divided into windows of 8 data points (∼20 μs), and the average value (dwell level) and the standard deviation (SD) of respective data windows were calculated. Dwells were distinguished from steps by a threshold value for SD. Data windows with an SD value of less than 1.5 were deemed to be dwells. The significance of differences between adjacent dwells was evaluated by Student’s *t*-test. Pairs without significant differences (*P* > 0.05) were concatenated, and step levels were updated with the mean of adjacent dwell levels. The statistical accuracy of the dwells determined by the *t*-test was further evaluated by examining whether the levels of adjacent dwells were outside the 3-sigma range of each other. These evaluations were repeated until all consecutive pairs of dwells were significantly distinguished. To determine slopes of the step periods, a sigmoid curve was fitted to a pair of step events, each of which consists of a slope and its preceding and succeeding dwells, using the dwell level as an initial value.

## Statistics and reproducibility

Statistical tests, sample size, and the number of biological replicates are given in the figure legends.

## Data availability

All data generated during this study are included in this published article, its Supplementary Information, and its Source Data file. *Salmonella* strains, a plasmid encoding pHluorin(M153R) and all other data are available from the corresponding author upon request.

## Acknowledgements

We acknowledge Michael D. Manson for critical reading of the manuscript and helpful comments and suggestions, and Yoshiyuki Sowa, Tomoko Miyata, Kazuki Kasai, and Fumiaki Makino for technical supports. We also thank Toshio Yanagida, Fumio Oosawa, Sho Asakura, Hirokazu Hotani, Donald L. D. Caspar, Seishi Kudo and Masahiro Ueda for encouragement, support, discussion, and critical reading of the manuscript. This work was supported by MEXT KAKENHI Grant Number JP25117501 to S.N., JP26115720 and JP15H01335 to Y.V.M., JP24117004, JP25121718, JP20H05532, and JP22H04844 to T.M. and JSPS KAKENHI Grant number JP24770141 to S.N., JP15H05593 and JP23H04082 to Y.V.M., JP19H03182, JP22H02573, and JP22K19274 to T.M. and JP25000013 to K.N., and JST PRESTO Grant Number JPMJPR204B to Y.V.M. This work has also been partially supported by JEOL YOKOGUSHI Research Alliance Laboratories of The University of Osaka to K.N.

## Author Contributions

S.N., T.M. and K.N. planned the project; S.N., Y.V.M. and N.K. carried out the experiments; N.K. set up the nanophotometry system including both hardware and software. S.N., Y.V.M. and K.N. analysed the data, and S.N., T.M. and K.N. wrote the paper based on discussion with other authors.

## Supplementary Information

**Extended Data Fig. 1.**
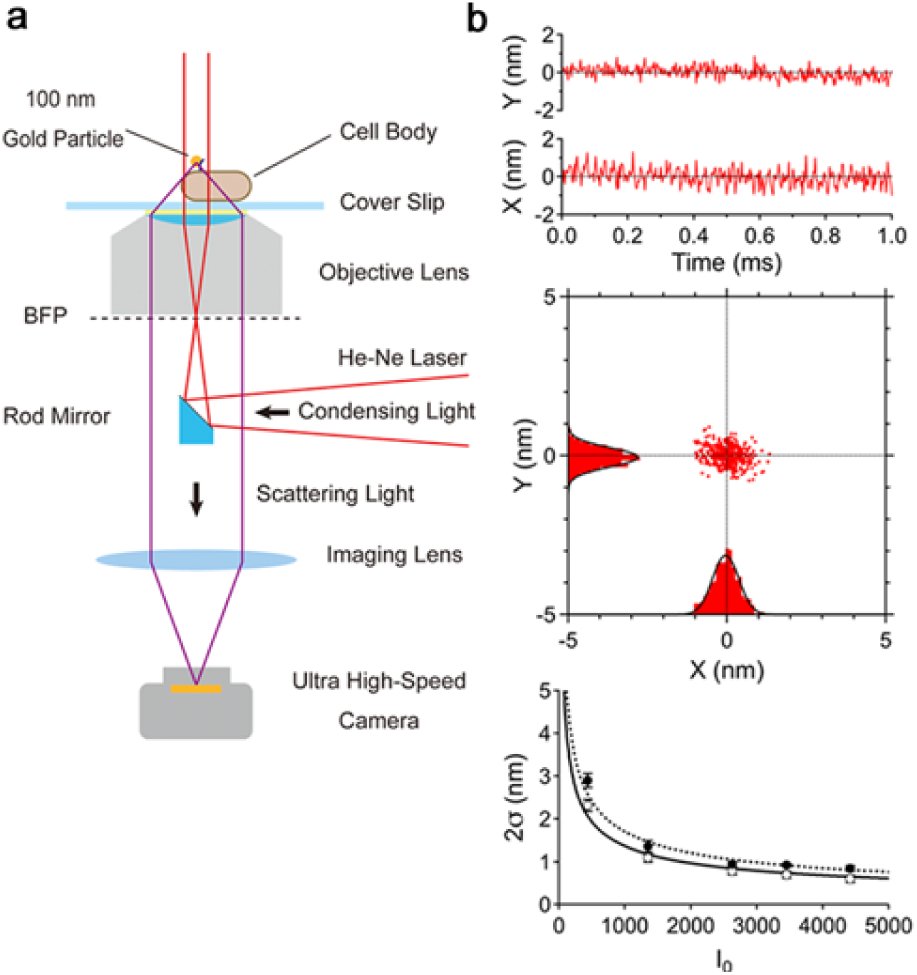
High-resolution measurement of flagellar motor rotation. (a) Schematic diagram of a back-scattering nanophotometry system used in this study. (b) Precision of the system in determining the position of a 100-nm AuNP firmly attached on a cover slip. Upper and middle panels show variations in X and Y positions at every 2.56-μs frame over 1 ms (392 data points each), and lower panel presents 2σ of Gaussian functions fitted to every 392 X and Y positions as a function of scattering light intensity. The smallest 2σ obtained at the highest intensity was 0.82 nm in X and 0.6 nm in Y. The standard deviations are plotted in the lower panel but difficult to see them because of their small sizes close to those of closed (X) and open (Y) circles.

**Extended Data Fig. 2.**
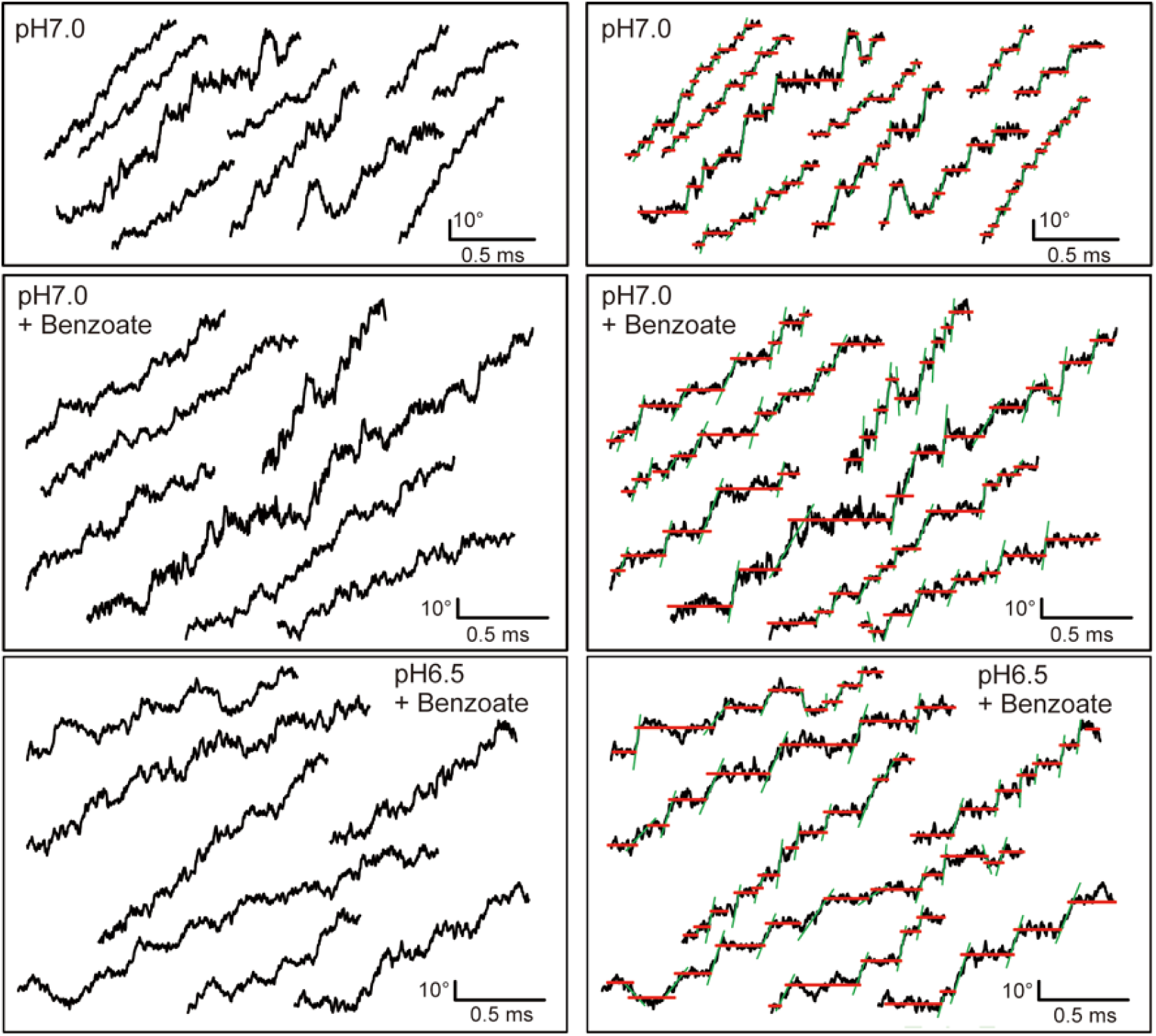
Typical steps extracted from the angle vs. time plots of the CCW-locked wild-type motor measured at an external pH value of 7.0 or 6.5 in the presence and absence of 5 mM potassium benzoate (left panels). The data of right panels are the same as those of left panels, and red and green lines are the results of fitting by the automated step finder.

**Extended Data Fig. 3.**
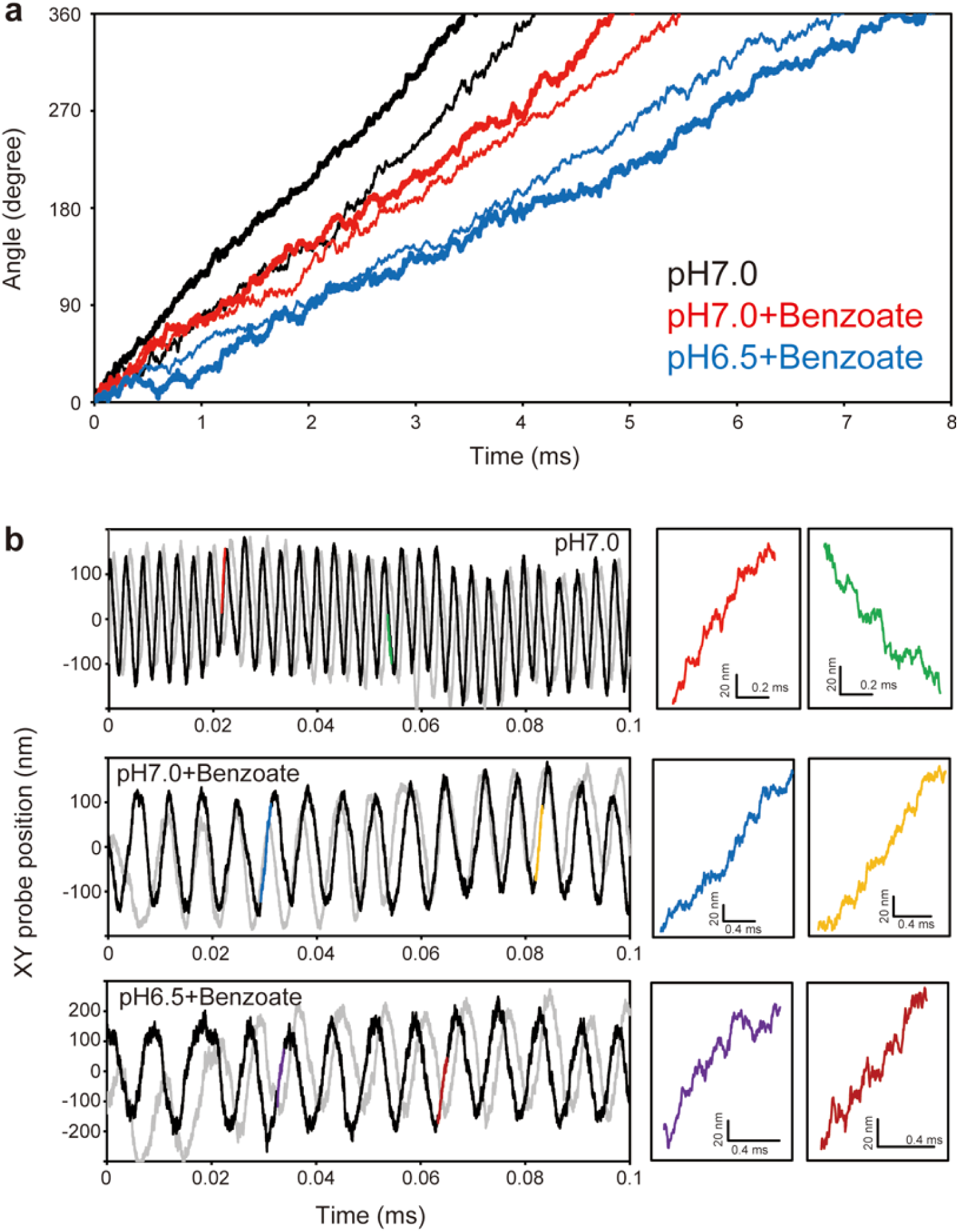
More examples of angular displacement of the CCW-locked wild-type motor measured in the presence and absence of 5 mM potassium benzoate. (a) Angle vs. time plots. Black, red, and blue data show angle traces measured in motility medim (pH 7.0) without benzoate, in motility medium (pH 7.0) containing benzoate, and in the motility medium (pH 6.5) containing benzoate, respectively. Thick and thin lines in each colour represent two distinct motors. (b) Time record of the probe position. Black and grey lines represent the X and Y displacements, respectively. Regions highlighted in colour are extended in left panels.

**Extended Data Fig. 4.**
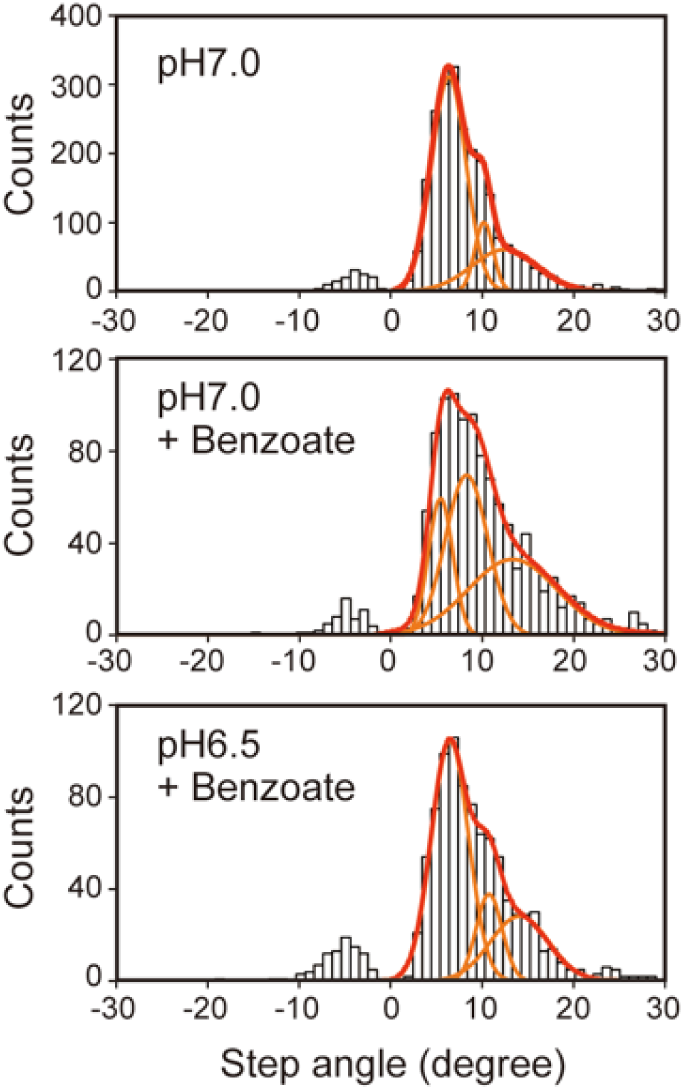
Step angles. Histograms of step angles were reproduced by the multi-Gaussian distribution (red lines) composed of three single-peak Gaussian distributions (orange lines). The multi-peak Gaussian fitting was performed using Igor Pro.

**Extended Data Fig. 5.**
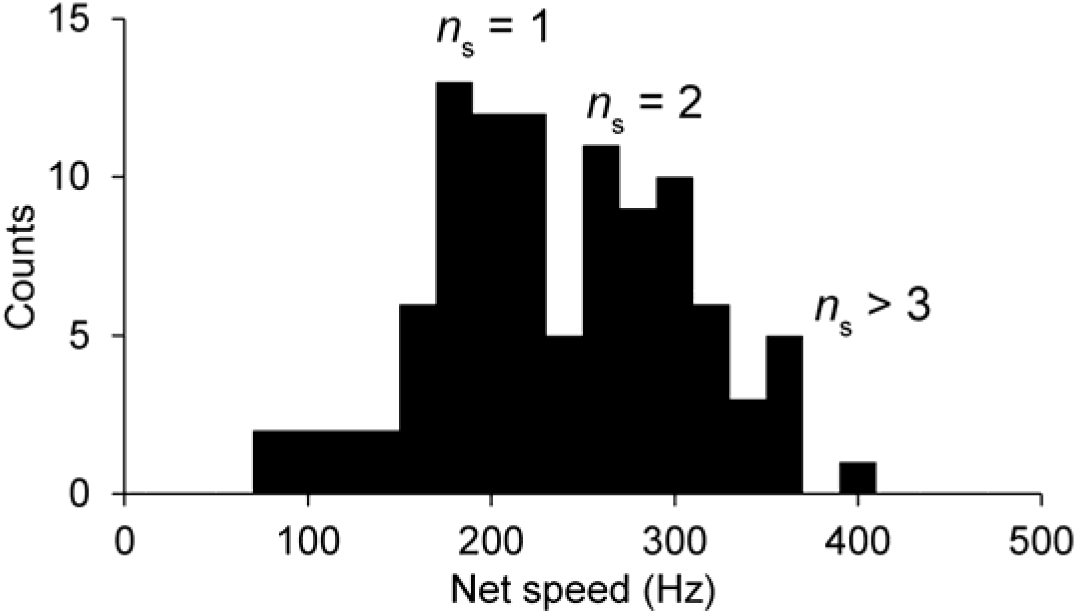
Net speed of the flagellar motor measured in this study. The net speeds were determined every one revolution by line fitting to rotation angle vs time plots. The number of stator units (*n*_s_) estimated from the speed distribution are shown.

**Extended Data Fig. 6.**
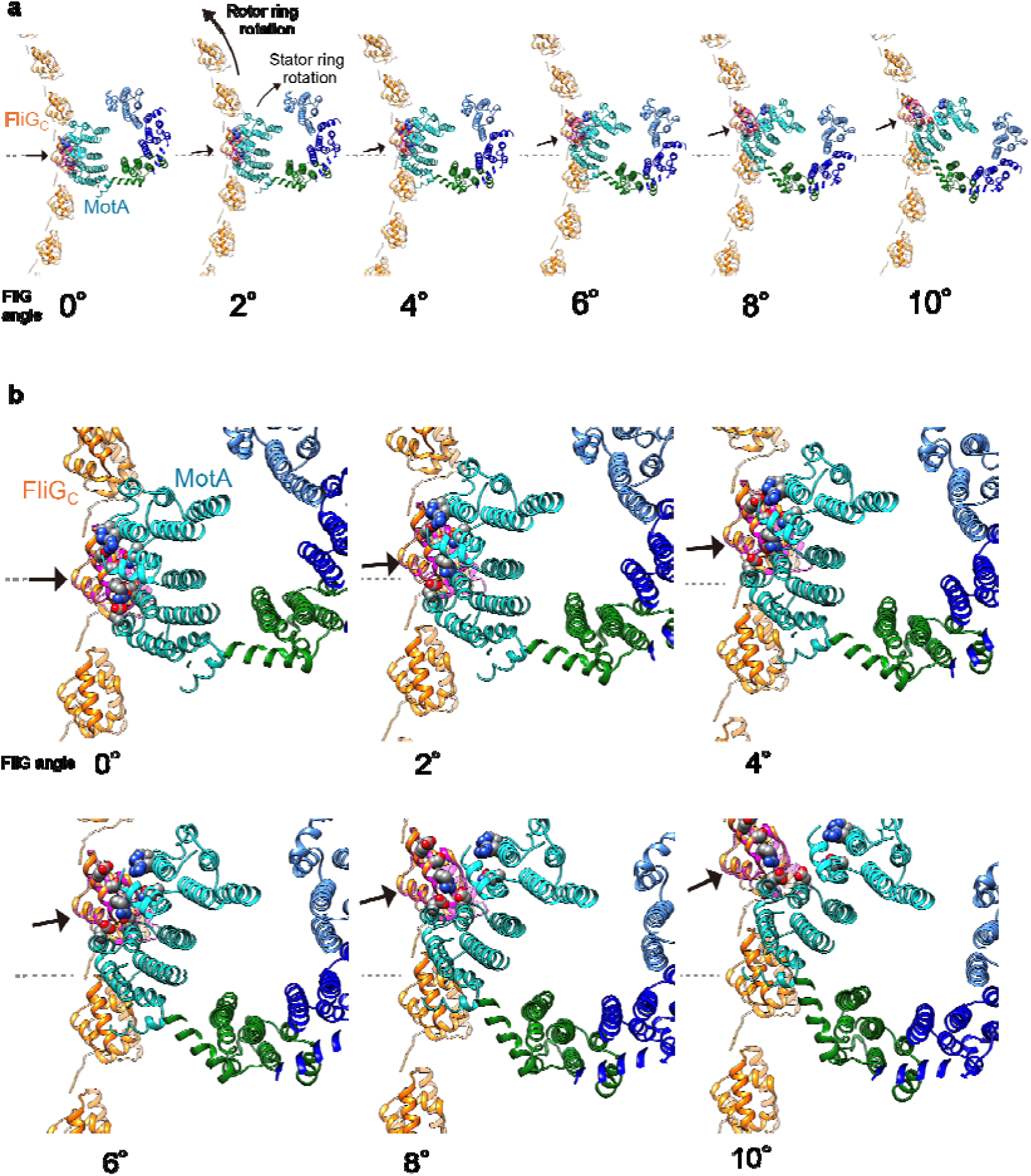
FliG_C_-MotA interaction cycle during rotation. Dissociation of FliG_C_ from MotA during the CW rotation of the MotA_5_ ring with the coupled CCW rotation of the FliG_C_ ring in the basal body C ring. The rotation of the FliG_C_ ring is depicted every 2° in both **a** and **b** where **b** is a magnified view of **a**. The tight bonding interactions of FliG_C_ and MotA start to be broken already at 2° rotation.

**Extended Data Fig. 7.**
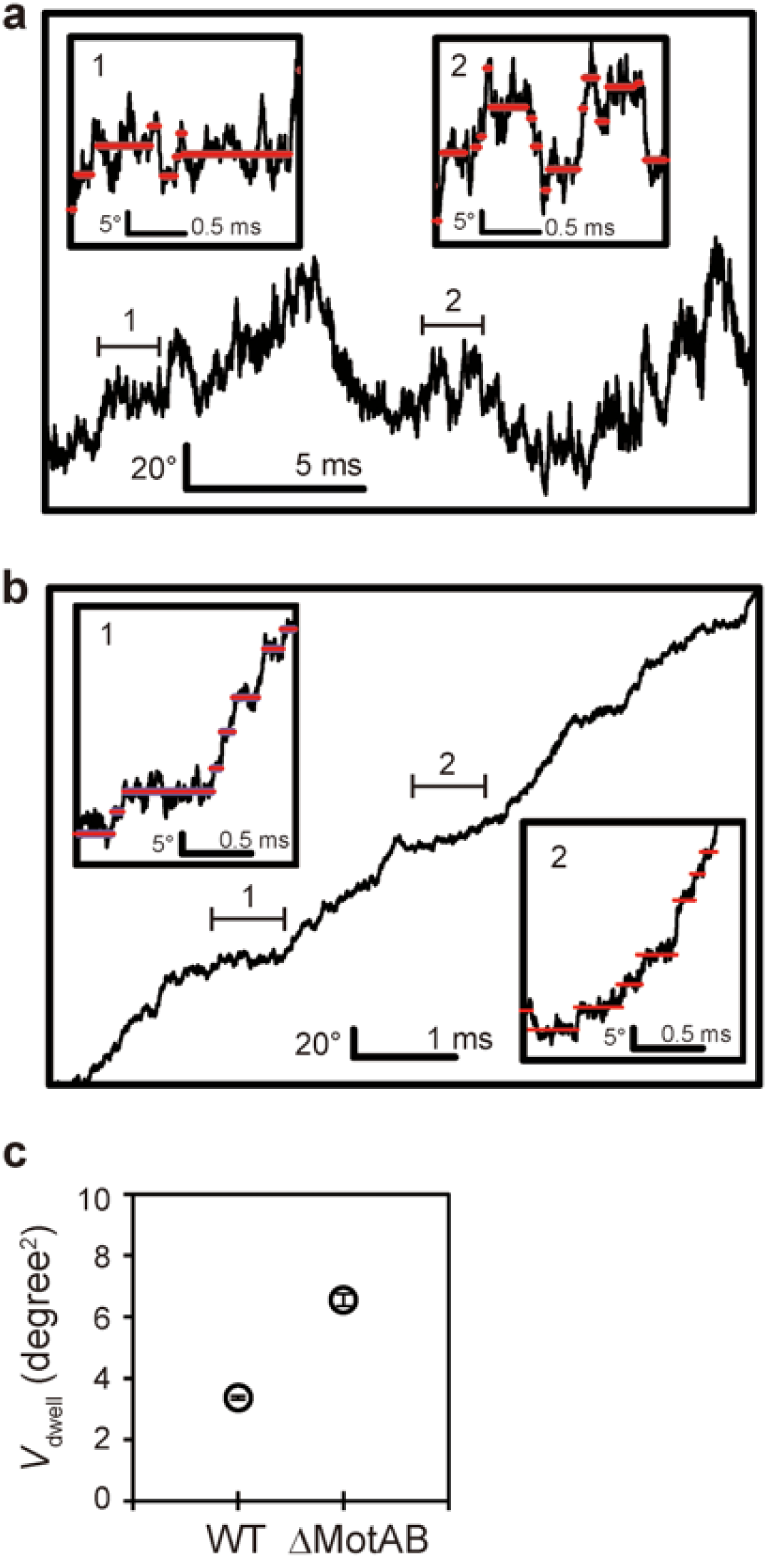
**Comparison of dwell times between the CCW-locked wild-type (WT) and stator-less mutant (**Δ**MotAB) motors.** Angle vs. time plots of **(a)** the ΔMotAB motor and **(b)** WT motor. The regions indicated with horizontal bars are extended in the insets. Red lines in the insets are dwell levels determined by the step finder. **(c)** Variance of the probe position in the dwell time. The average values and standard errors are shown.

---

**Extended data Table 1.**

| | Peak angle $\pm \sigma^*$ | | |
| --- | --- | --- | --- |
|  | pH 7.0 | pH 7.0 | pH 6.5 |
|  | without benzoate | +5mM benzoate | +5mM benzoate |
| Peak 1 | $6.3 \pm 1.9^\circ$ | $5.4 \pm 1.3^\circ$ | $6.5 \pm 2.0^\circ$ |
| Peak 2 | $10.1 \pm 1.0^\circ$ | $8.4 \pm 2.3^\circ$ | $10.8 \pm 1.4^\circ$ |
| Peak 3 | $12.5 \pm 3.7^\circ$ | $13.5 \pm 4.9^\circ$ | $14.2 \pm 3.2^\circ$ |

\*Peak angles and $\sigma$ values were determined by fitting the multi-peak Gaussian distribution to the step angle distributions.

**Extended Data Table 2.**

| Parameter |  | Value |
| --- | --- | --- |
| Periplasmic pH | $\text{pH}^{\text{Peri}}$ | 7.0 |
| Cytoplasmic pH | $\text{pH}^{\text{Cyto}}$ | 7.4 |
| Membrane potential | $\Delta\psi$ (mV) | 150 |
| Temperature | $T$ (K) | 296 |
| | $k_0$ ($\text{s}^{-1}$) | $7.0 \times 10^7$ |
| Periplasmic $\text{H}^+$ binding rate | $k_{\text{on}}^{\text{Peri}}$ ($\text{M}^{-1} \text{s}^{-1}$) | $5.0 \times 10^9$ |
| Periplasmic $\text{H}^+$ dissociation rate | $k_{\text{off}}^{\text{Peri}}$ ($\text{s}^{-1}$) | $3.0 \times 10^5$ |
| Cytoplasmic $\text{H}^+$ binding rate | $k_{\text{on}}^{\text{Cyto}}$ ($\text{M}^{-1} \text{s}^{-1}$) | $2.5 \times 10^8$ |
| Cytoplasmic $\text{H}^+$ dissociation rate | $k_{\text{off}}^{\text{Cyto}}$ ($\text{s}^{-1}$) | $1.5 \times 10^4$ |
| Dissociation constant | $k_{\text{off}}/k_{\text{on}}$ (M) | $6.0 \times 10^{-5}$ |
| Apparent periplasmic reaction rate* | $k_{\text{H}}^{\text{Peri}}$ ($\text{s}^{-1}$) | $3.0 \times 10^5$ |
| Apparent cytoplasmic reaction rate* | $k_{\text{H}}^{\text{Cyto}}$ ($\text{s}^{-1}$) | $1.5 \times 10^4$ |
| Power-stroke angle | $\theta_{\text{ps}}$ (degree) | 2 |

\*Apparent reaction rates were determined as follows: The reaction $S + \text{H}^+ \rightleftharpoons \text{SH}^+$ , where $S$ is a stator unit, with the protonation rate $k_{\text{on}}$ and deprotonation rates $k_{\text{off}}$ gives $[\dot{S}H] =$ $k_{\text{on}}[S][\text{H}^+] - k_{\text{off}}[\text{SH}^+]$ . Assuming $[\text{H}^+] \gg [S]$ and the initial concentrations of $S$ and $\text{H}^+$ as $[S]_0$ and $[\text{H}^+]_0$ , respectively, $[\text{H}^+]_0 - [\text{SH}^+] \sim [\text{H}^+]_0$ and then $[\dot{S}H] \sim k_{\text{on}}[S]_0[\text{H}^+]_0 -$ $(k_{\text{on}}[\text{H}^+]_0 + k_{\text{off}})[\text{SH}^+]$ . Because $[S]_0[\text{H}^+]_0$ is constant, $k_{\text{on}}[\text{H}^+]_0 + k_{\text{off}}$ was used as the apparent reaction constant.

